# MicroRNA-22, Specificity protein-1 and Cystathionine β-synthase in early onset Preeclampsia: significance in trophoblast invasion

**DOI:** 10.1101/2023.03.08.531738

**Authors:** Pallavi Arora, Sankat Mochan, Sunil Kumar Gupta, Neerja Rani, Pallavi Kshetrapal, Sadanand Dwivedi, Neerja Bhatla, Renu Dhingra

## Abstract

**Introduction:** Trophoblast cell invasion during human placentation is majorly regulated by the balance between MMPs 2, 9 and their inhibitors [tissue inhibitors of matrix metalloproteinases (TIMPs 1, 2)]. Exogenous NaHS (hydrogen sulphide donor) treatment was shown to significantly increase the expression levels of matrix metalloproteinases (MMPs 2, 9) in human bladder cancer EJ cells. Epigentically, the gene expression of hydrogen sulphide synthesising enzyme CBS (cystathionine β-synthase) could be further regulated by various mi-RNAs via the transcription factors like Sp1. Specificity protein 1 (Sp1) has been identified as a target gene for miR-22 to regulate the invasion and metastasis of gastric cancer cells. However, the mechanism of MMPs regulation by either CBS, Sp1 and miRNA-22 in the pregnancies having early onset preeclampsia is not known.

**Aim of the study:** To determine and compare the expression of MMPs 2, 9, TIMPs 1, 2, CBS, Sp-1 and miR-22 in early onset preeclamptic patients and normotensive, non-proteinuric controls at both transcription and translation levels.

**Materials and methods:** 30 pregnant women were enrolled from Department of Obstetrics and Gynaecology, AIIMS, New Delhi, India. EOPE women (n=15) after clinical diagnosis as per ACOG guidelines were enrolled as cases and normotensive, non-proteinuric pregnant women (n=15) were enrolled as controls. Protocol of the study was approved by Institute Ethics Committee, AIIMS, New Delhi. 30 caesarean delivered placentae (15 each of patients and controls) were collected to analyse the mRNA and protein expression/levels of MMPs 2, 9, TIMPs 1, 2, CBS, and Sp1. Protein activity of MMP-2 and 9 was determined. Gene expression of miR-22 was observed in placentae of the recruited patients and controls. Data were analysed by STATA 14 and Graph Pad Prism 8.

**Results:** Significantly reduced mRNA and protein expression/levels of MMPs 2, 9, CBS and Sp1 whereas elevated for those of TIMP-1 and TIMP-2 was observed in EOPE patients as compared to controls. Gelatinolytic activity of MMP-2 and 9 was also found significantly reduced in placentae from EOPE patients whereas gene expression of miR-22 was found significantly up regulated in the early onset preeclamptic patients in comparison to controls.

**Conclusion:** This is the first study of its kind which implicates that insufficient trophoblastic invasion may be because of down regulation of MMPs 2, 9, CBS, Sp1 and concomitant up regulation of TIMPs 1, 2 and miR-22 in the early onset preeclamptic patients as compared to controls.

## Introduction

Preeclampsia (PE) is a multisystem disorder of pregnancy defined by high blood pressure and proteinuria^1^. The hallmark of PE is the impaired capacity of the trophoblast to invade the uterine spiral arteries, which results in a poorly perfused fetoplacental unit^2-4^. Depending on time, PE is classified as early-onset preeclampsia, which requires delivery before 34 weeks of gestation, or late-onset preeclampsia, with delivery at or after 34 weeks or later^5-9^. It is thought that early onset PE poses a high risk to both mother and fetus^10,11^ whereas late onset PE may present with less severe clinical symptoms^12^. Impaired trophoblast invasion observed in early stage of PE is characterized by a decrease in MMP-2 and MMP-9, which affects spiral artery remodelling, causing a dysfunctional uteroplacental circulation^13^. MMPs and their inhibitors (TIMPs) play a major role in trophoblast invasion into the uterine wall^13^. The identification of changes in the levels and activity of MMPs 2, 9 as well as their endogenous inhibitors (TIMPs 2, 1) in both defective trophoblast invasion and endothelial dysfunction led to the consideration of these proteases as key mediators in the pathological features of preeclampsia^13^. Hydrogen sulphide (H_2_S) has been reported to influence MMP-2 expression in-vitro, they treated hepatocellular carcinoma or hepatoma cells with NaHS (H_2_S donor) and observed up regulated MMP-2 protein expression suggesting hydrogen sulphide promotes invasion process in various cancer cell lines^14,15^. Endogenously, H_2_S is produced as a metabolite of homocysteine (Hcy) by cystathionine β-synthase (CBS), cystathionine γ-lyase (CSE), and 3-mercaptopyruvate sulfurtransferase (3MST)^16^. The regulation of CBS -1b and -1a promoters in a proliferation-sensitive manner by specificity protein 1 (Sp1) has already been reported^17^, in the study, the authors have transfected Sp1 deficient fibroblasts cells by Sp1 expression construct and observed significantly high levels of CBS expression^17^. Sp1 was identified as a novel, direct target of miRNA-22 using luciferase reporter assay^18^. miRNA-22 overexpression diminished but miRNA-22 knockdown up regulated Sp1 mRNA and protein expression in colorectal carcinoma cells suggesting an inverse relation between miRNA-22 and Sp1^18^. Also, miR-22 expression was found to be increased in EOPE placentae as compared to placentae from preterm labor controls^19^.

The aim of the present study was to determine the expression/levels of MMPs 2, 9, TIMPs 1, 2, CBS, Sp1 and miRNA-22 in the pregnancies complicated by early onset preeclampsia and the comparison was done with those of controls.

## Materials and methods

### Study Subjects

30 pregnant women were enrolled from the antenatal clinic and the ward of the Department of Obstetrics and Gynaecology, All India Institute of Medical Science, New Delhi, India. Early onset preeclamptic women (n=15) after clinical diagnosis as per ACOG guidelines were enrolled as cases and normotensive, non-proteinuric pregnant women (n=15) (maternal and gestational age matched) were enrolled as controls. Protocol of the study was approved by the Institute Ethics Committee, AIIMS, New Delhi (Ref. No. IECPG-247/30.03.2016) and written informed consent was obtained from all the enrolled women. 15 each of early onset PE patients and maternal age matched control placentae (caesarean delivered) were collected.

Placental samples were used subsequently to determine and compare the gene expression of MMPs 2, 9, TIMPs 1, 2, cystathionine beta synthase, specificity protein-1, precursor miR-22 and miR-22-3p in early onset preeclamptic patients and normotensive, non-proteinuric pregnant women by qRT-PCR (quantitative Real Time-Polymerase Chain Reaction).

### quantitative Real Time-Polymerase Chain Reaction

Placental tissue (kept in RNA later) was blotted on absorbent paper and 100 mg was taken, washed in 1× 0.1 M PBS followed by RNA isolation using Ambion, Invitrogen. The quality of RNA from placentae was examined by denaturing gel and quantity was measured on Micro-Volume UV/Visible Spectrophotometer (Thermo Fisher Scientific-NanoDrop TM 2000). c-DNA preparation was done using Revert aid H-minus reverse transcriptase kit (Thermo). Quality of cDNA was checked on 0.8% agarose gel, visualized by ethidium bromide (EtBr) stain under UV and quantity was measured on Micro-Volume UV/Visible Spectrophotometer (Thermo Fisher Scientific-NanoDrop TM 2000). cDNA was amplified by quantitative RT-PCR (CFX96 Touch™ Real-Time PCR Detection System, BioRad). qRT-PCR reactions were carried out in 20 µl volume, including cDNA (template), SYBR Green (Thermo), forward and reverse primers (Sigma), and nuclease free water to determine mRNA expression of MMPs 2,9, TIMPs 1,2, CBS, Sp1 and gene expression of miR-22 (pre miR-22 and miR-22-3p). Glyceraldehyde-3-phosphate dehydrogenase (GAPDH) mRNA and U6 small nuclear RNA were used as internal controls. Primers were designed by NCBI and confirmed by In-Silico PCR (Table 1).

**Table 1:**
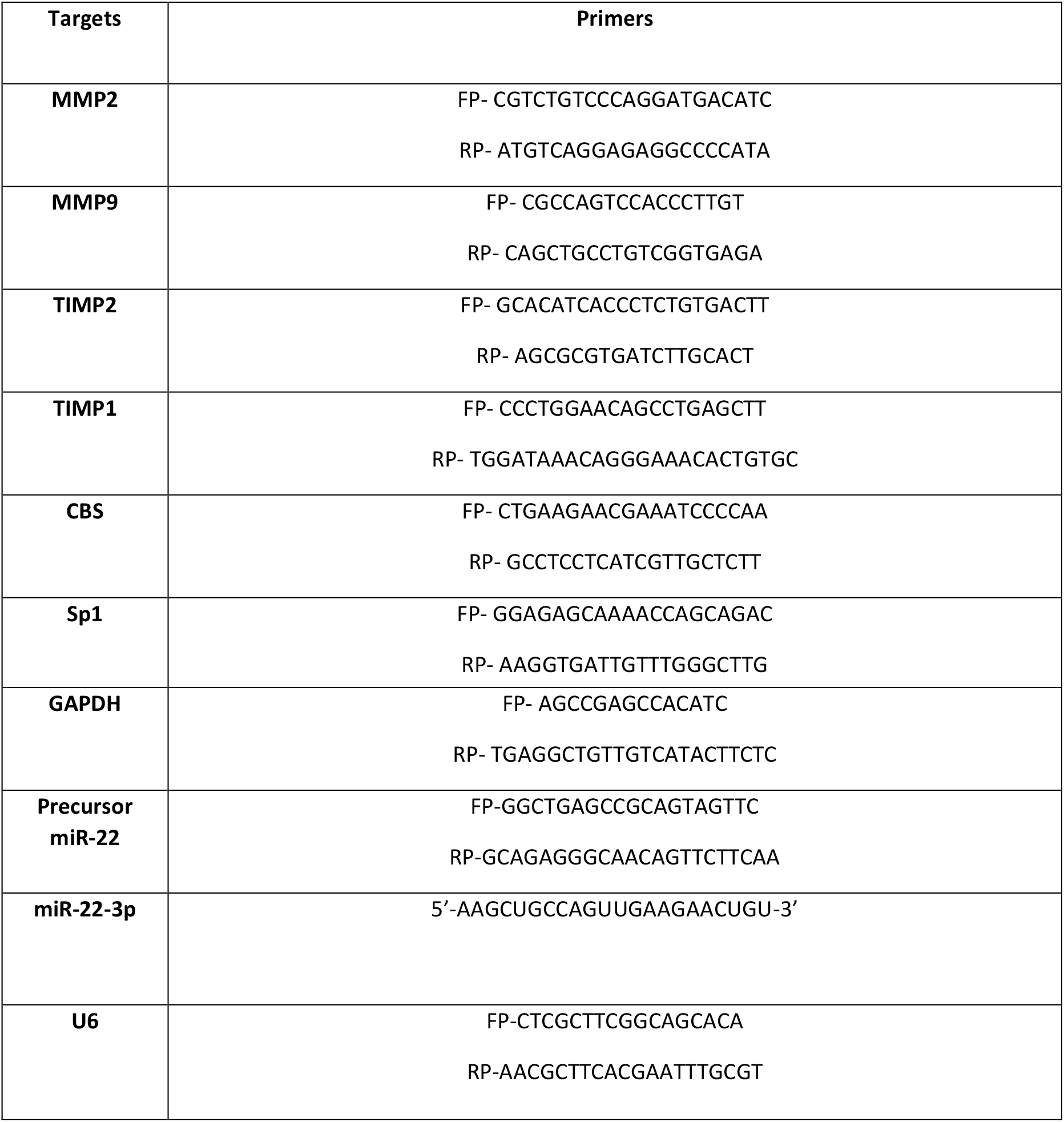
Primers designed by NCBI (National Centre for Biotechnology Information) and confirmed by In-Silico PCR. GAPDH: Glyceraldehyde 3-phosphate dehydrogenase.

The protein expression of the above-mentioned markers was observed by immunohistochemistry (IHC) and immunofluorescence (IF) and their levels were determined by western blot in the placentae of early onset preeclamptic patients and normotensive, non-proteinuric controls. In all the collected placentae, protein activity of MMP-2 and MMP-9 was seen by gelatin gel zymography.

### Immunohistochemistry

Paraffin tissue blocks were sectioned on microtome (Thermo Scientific™ HM 325) (5µm sections) and were taken on Poly-L-lysine (Sigma) coated slides. UltraVision™ Quanto Detection System HRP DAB (Thermo, TL-125-QHD) was used to determine protein expression of MMPs 2,9, TIMPs 1,2, CBS and Sp1. Primary antibodies to MMP-2 (Abcam) at a dilution of 1:500, MMP-9 (Abcam) at a dilution of 1:500, TIMP-1 (Thermo) at a dilution of 1:100, TIMP-2 (Novus Biologicals) at a dilution of 1:100, CBS (Abcam) at a dilution of 1:400 and Sp1 (Merck) at a dilution of 1:250 were used. Slides were counterstained with hematoxylin, dehydrated in graded ethanol, cleared, and cover slips were applied. Mounting was done by Dibutylphthalate Polystyrene Xylene (DPX). Stained slides were observed under Nikon Eclipse Ti-S elements microscope using NiS-AR software (version 5.1). Other chemicals were procured from Fisher Scientific.

### Immunofluorescence

Paraffin tissue blocks were sectioned on microtome (Thermo Scientific™ HM 325) and were taken on Poly-L-lysine (Sigma) coated slides. Two changes of xylene (each for 5 minutes) followed by two changes of absolute alcohol (each for 3 minutes) and subsequently one change of 90% alcohol (for 1 minute) were given. Slides were rinsed with distilled water. Antigen retrieval was done with sodium citrate buffer (15 minutes at 95-100°C) followed by treatment with TBST× (2 minutes) (2 times). BSA (blocking agent) was applied on slides for 35 minutes followed by overnight incubation with primary antibodies [MMP-2 (Abcam) at a dilution of 1:100, MMP-9 (Abcam) at a dilution of 1:100, TIMP-1 (Thermo) at a dilution of 1:10, TIMP-2 (Novus Biologicals) at a dilution of 1:25, CBS (Abcam) at a dilution of 1:200 and Sp1 (Merck) at a dilution of 1:50] at 4^0^C. Slides were then rinsed with PBST20 followed by incubation with secondary antibodies [MMP-2, MMP-9, TIMP-2, CBS, Sp1 (FITC conjugated, Abcam), TIMP 1 (TRITC conjugated, Abcam)] for 2.5 hours and then washed with PBS. Mounting was done by fluoroshield mounting media with DAPI (Abcam). Stained slides were observed under Nikon Eclipse Ti-S elements microscope using NiS-AR software (version 5.1). Other chemicals were procured from Fisher Scientific.

### Western Blot

Protein extraction (from placental tissues of both patients and controls) was done with RIPA buffer (Thermo) and protease inhibitor cocktail. Separating and stacking gels were prepared. 3× non-reducing sample buffer was added to each of isolated protein sample from patients and controls followed by denaturation at 95 ° C for 5 minutes and the samples were loaded along with protein molecular marker to the wells. Subsequently, running the gel at 50 V (vertical electrophoresis apparatus, BioRad) in electrophoresis buffer for 3-3.5 hours until good band separation is achieved. Nitrocellulose membrane was used for the transfer of gel products on to the membrane in transfer buffer for 90 minutes. Membrane was washed with TBST20 followed by blocking in 5% BSA TBST20 for 90 minutes. Overnight incubation was done with primary antibodies [MMP-2 (Abcam) at a dilution of 1:1000, MMP-9 (Abcam) at a dilution of 1:1000, TIMP-1 (Thermo) at a dilution of 1:200, TIMP-2 (Novus Biologicals) at a dilution of 1:200, CBS (Abcam) at a dilution of 1:1000 and Sp1 (Merck) at a dilution of 1:400] at 4^0^C. Washing was done with TBST20 followed by incubation with secondary antibodies (Abcam) for 3 hours and then washed with TBST20. ECL kit (Thermo) was used for visualization of bands in Densitometer (Protein Simple).

### Gelatin gel zymography

Protein extraction (from placentae of both patients and controls) was done with RIPA buffer (Thermo) and protease inhibitor cocktail. 7.5% acrylamide gel was prepared containing gelatin. 5× non-reducing sample buffer was added to each of isolated protein sample from patients and controls. 10 μl protein sample was loaded to each well. Protein molecular weight marker (Thermo) was loaded, subsequently, running the gel at 90 V (vertical electrophoresis apparatus, BioRad) in electrophoresis buffer until good band separation was achieved. Gel was washed (2 × 30 minutes) with washing buffer then rinsing in incubation buffer for 5–10 minutes at 37°C with agitation. Fresh incubation buffer was added to the gel followed by incubation at 37°C for 24 h. Gel was stained with Coomassie blue for 30 minutes and rinsed with water. Incubation was done with destaining solution to visualise the bands showing pro and active forms of MMP-2 and MMP-9 in the study groups.

### Statistical Analysis

Data was analyzed by STATA 14 and Graph Pad Prism 8. Relative quantification cycles of gene of interest (ΔCq) were calculated by ΔCq = Cq (target) - Cq (reference). Relative mRNA expression with respect to internal control gene was calculated by 2^-ΔCq^. Paired t-test and Wilcoxon matched-pairs signed rank test were used to compare the average level of the variable between two groups, *p* value<0.05 was considered statistically significant.

## Results

Clinical characteristics of early onset preeclamptic patients and their gestational and maternal age matched normotensive, non-proteinuric controls are mentioned in Table 2.

**Table 2:** Maternal study population - clinical characteristics, n= number of subjects, Data presented as mean±SD, *p* values between groups were evaluated by Paired t-test, *p*<0.05 considered statistically significant.

| Study Groups |  |  |  |
| --- | --- | --- | --- |
| Clinical characteristics | EOPE (n=15) | Normotensive, Non proteinuric (Controls) (n=15) | Paired t test (*p* value) |
| Maternal age (years) | 29 ± 3.94 | 29.9 ± 4.44 | *p*>0.05 |
| Gestational age (onset) (weeks) | 30.68 ± 2.88 | 31.32 ± 3.65 | *p*>0.05 |
| Gestational Age (delivery) (weeks) | 34.26 ± 4.20 | 37.57 ± 1.26 | 0.002* |
| Systolic blood pressure (mmHg) | 146.82 ± 10.59 | 115.26 ± 7.97 | *p*<0.0001* |
| Diastolic blood pressure (mmHg) | 95.8 ± 8.97 | 74.72 ± 7.51 | *p*<0.0001* |
| Protein (dipstick) | 27- 1+, 13- 2+, 10- 3+ | Nil or traces | Not applicable |
| Placental weight | 440 ± 47.71 | 507.8 ± 9.60 | 0.04* |

### Significantly down regulated expression of MMPs 2, 9 whereas elevated for those of TIMPs 1, 2 in the early onset preeclamptic patients

qRT-PCR data revealed significantly reduced mRNA expression of MMPs 2, 9 whereas upheaved for those of TIMPs 1, 2 in the placental samples from pregnancies complicated by early onset preeclampsia as compared to the normotensive, non-proteinuric controls [Figure 1]. Immunostaining demonstrated weaker expression of MMP-2 and MMP-9 in syncytiotrophoblasts, stroma and around blood vessels in the placentae from early onset PE patients as compared to controls [Figure 2,3]. TIMPs 1, 2 were found strongly localized in the stromal component and syncytium in the placentae from early onset PE patients as compared to control placentae [Figure 2,3]. Immunoblot data showed significantly down regulated expression of MMP s 2, 9 whereas up regulated for those of TIMPs 1, 2 in the placentae from early onset PE patients as compared to normotensive, non-proteinuric control placentae [Figure 4]. MMPs 2, 9 activity (both pro and active forms) was significantly lower in placentae from early onset preeclamptic patients as compared to placentae from controls [Figure 5,6].

**Figure 1.**
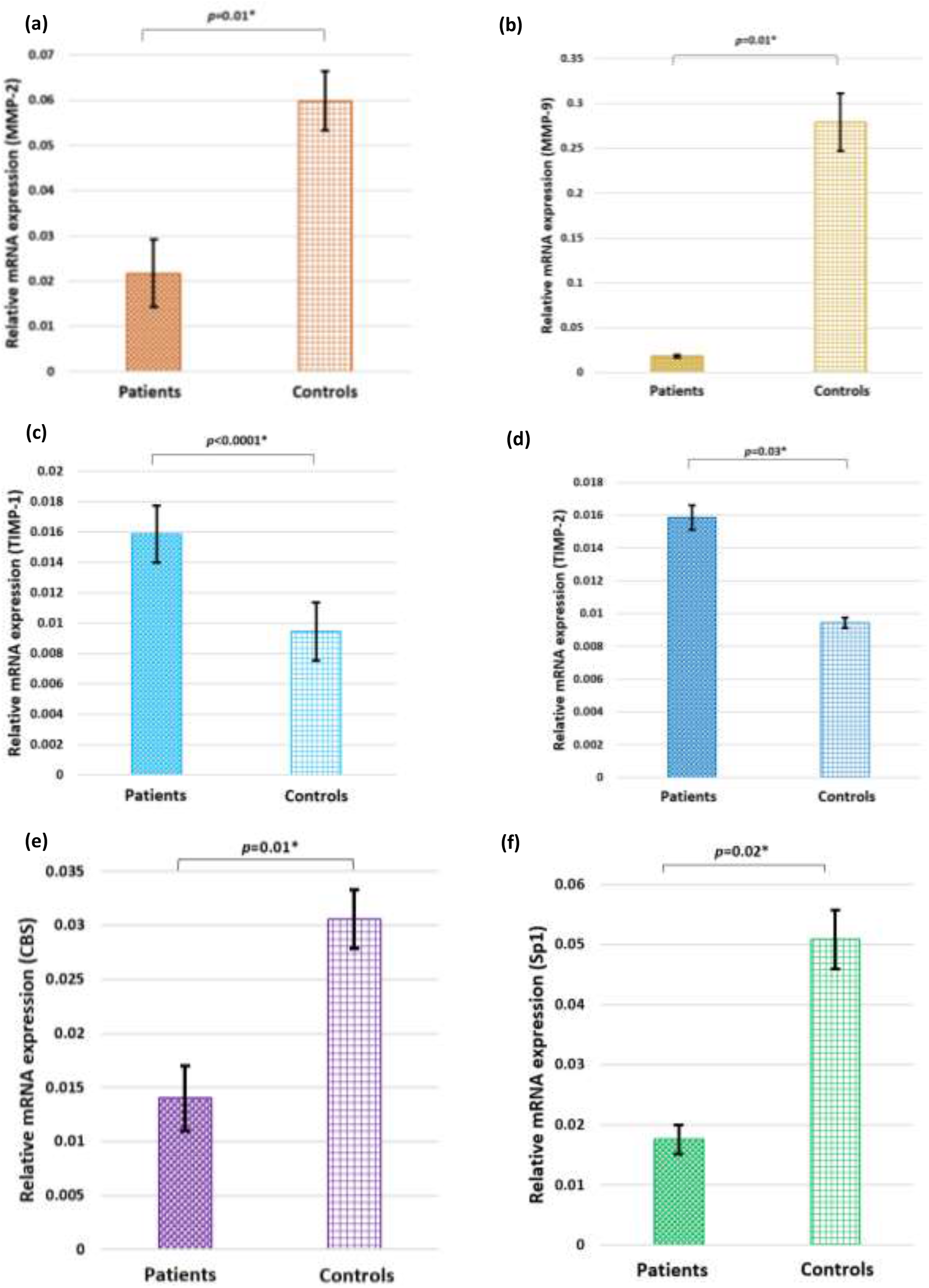
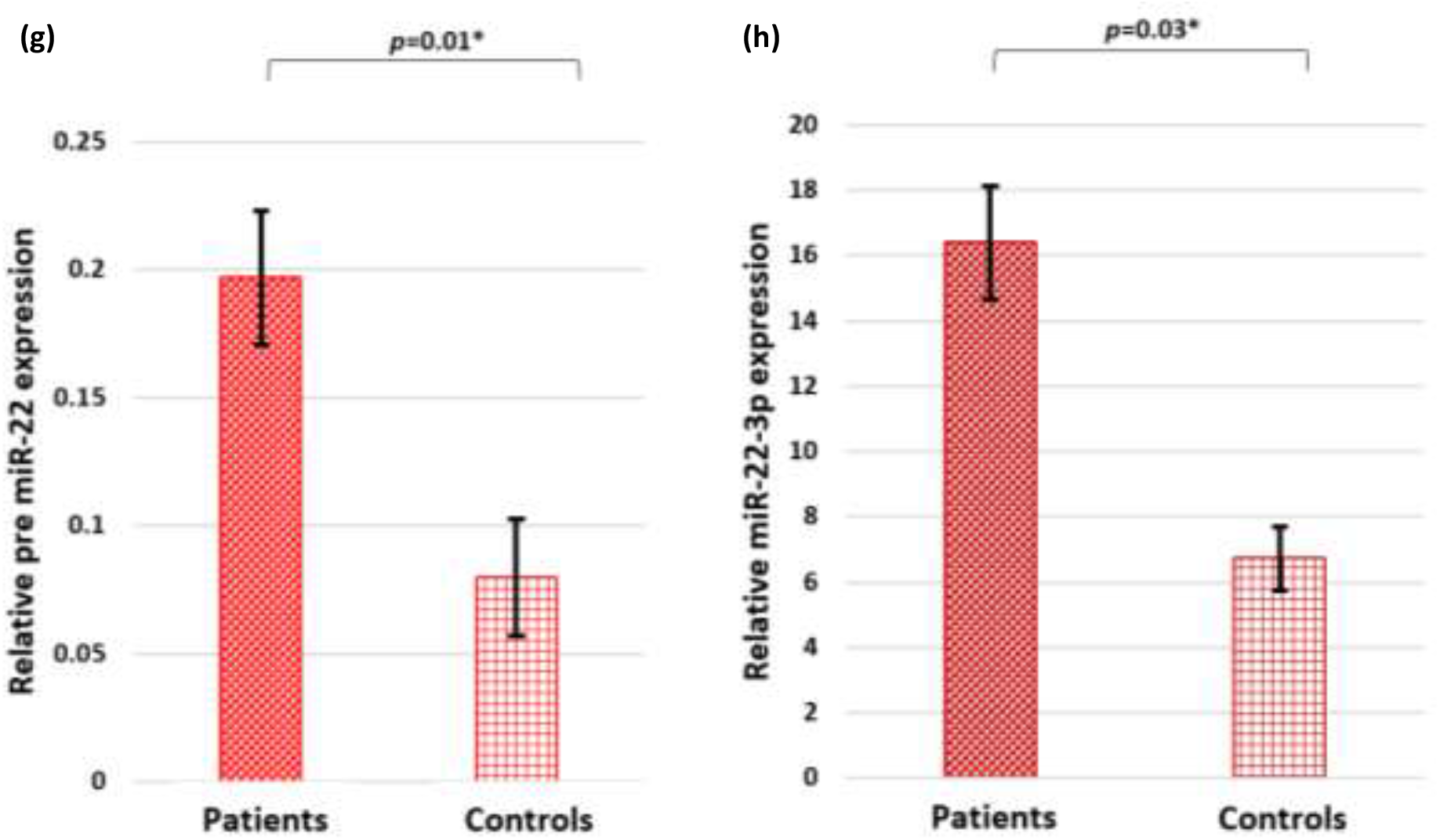
(a-f): Bar diagrams represent the relative mRNA expression of MMPs 2, 9, TIMPs 1,2, CBS and Sp1 in placentae of early onset preeclamptic patients and normotensive, non-proteinuric controls. GAPDH was used as positive control; g, h: Bar diagrams represent the relative precursor miR-22 and miR-22-3p expression. U6 was used as positive control. Data presented as mean ± SEM. Wilcoxon matched-pairs signed rank (MMPs 2, 9, TIMP-1, CBS, Sp1, pre miR-22, miR-22-3p) and paired t (TIMP-2) tests were applied, *p* values indicated on graphs.

**Figure 2:**
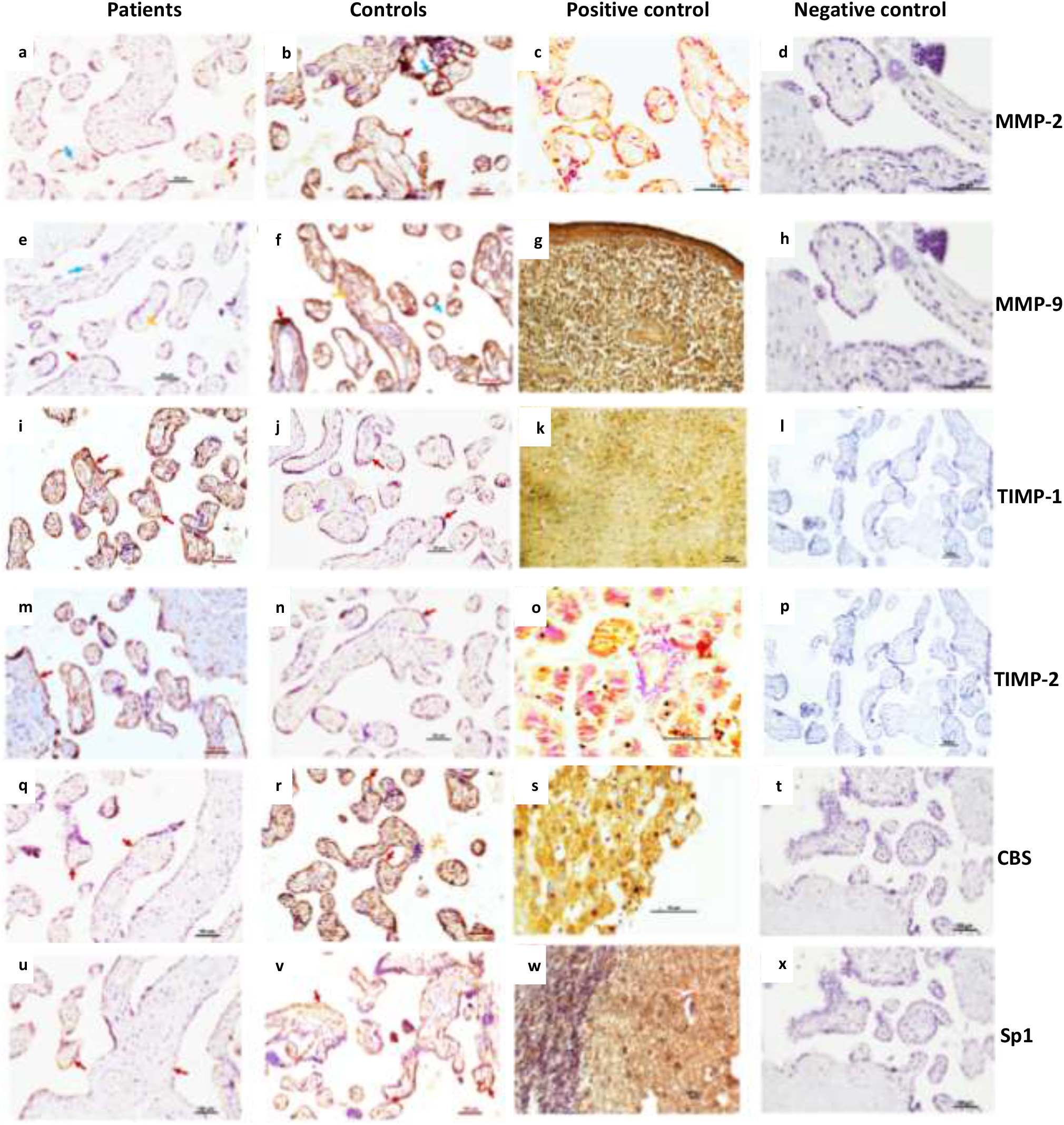
Representative immunohistochemistry images of placentae from early onset preeclamptic patients (a,e,i,m,q,u) and normotensive, non-proteinuric controls (b,f,j,n,r,v) showing MMP-2 (a-c), MMP-9 (e-g), TIMP-1 (i-k), TIMP-2 (m-o), CBS (q-s) and Sp1 (u-w) localization in syncytiotrophoblasts 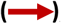, stromal component 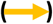 and blood vessels 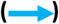. Positive controls: human placenta [MMP-2 (c)], human spleen [MMP-9 (g)], rat brain [TIMP-1 (k)], human pancreas [TIMP-2 (o)], human liver [CBS (s)], human cerebellum [Sp1 (w)].

**Figure 3:**
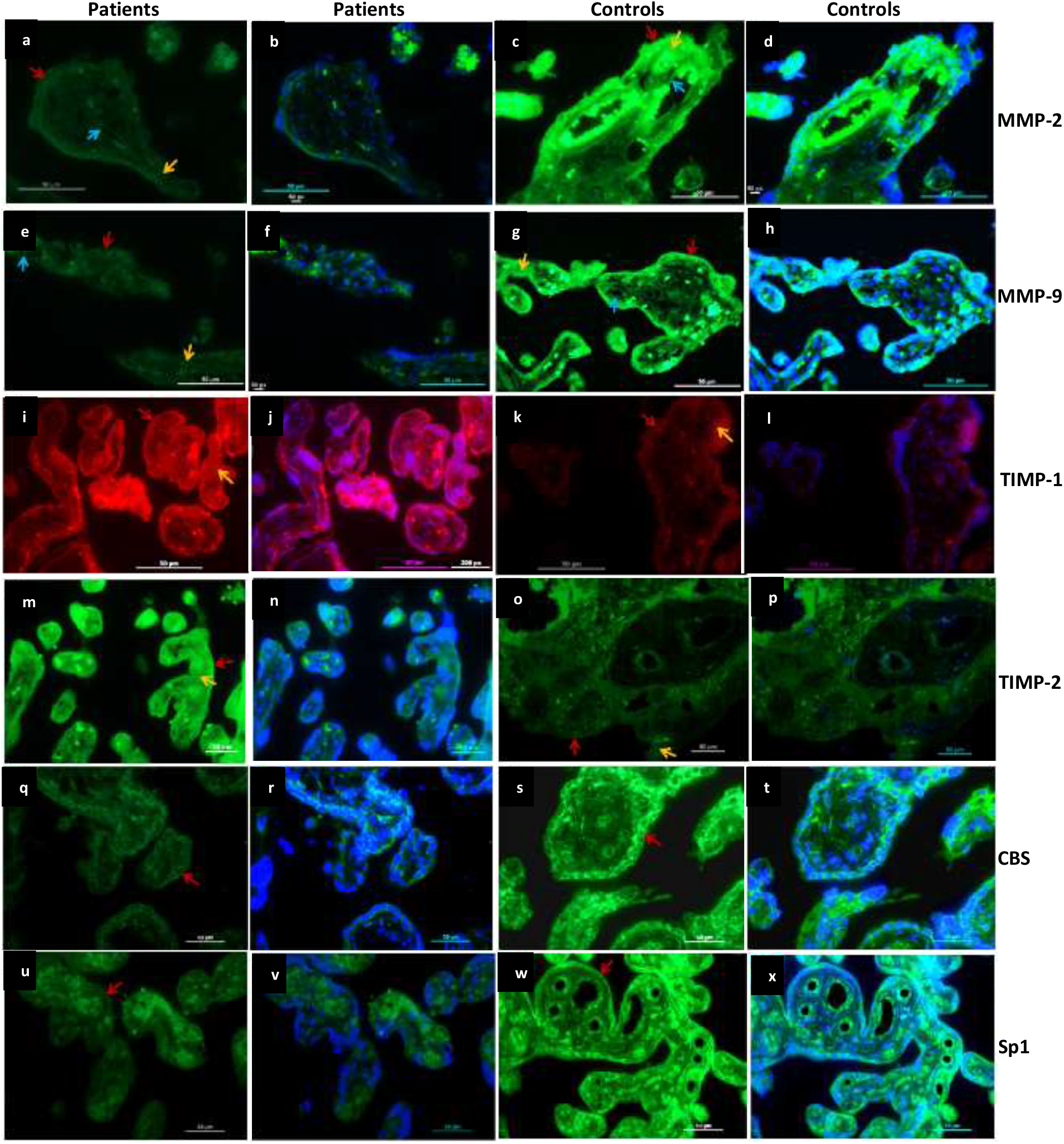
Representative Immunofluorescence images of placentae from early onset preeclamptic patients and normotensive, non-proteinuric controls showing MMP-2 (a-d), MMP-9 (e-h), TIMP-1 (i-l), TIMP-2 (m-p), CBS (q-t) and Sp1 (u-x) localization [MMPs 2, 9, TIMP-2, CBS, Sp1 (FITC), TIMP-1 (TRITC)] in syncytiotrophoblasts 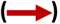, stromal component 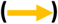 and blood vessels 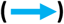. Nuclei were stained by DAPI; merged images: d,h,l,p,t,x

**Figure 4:**
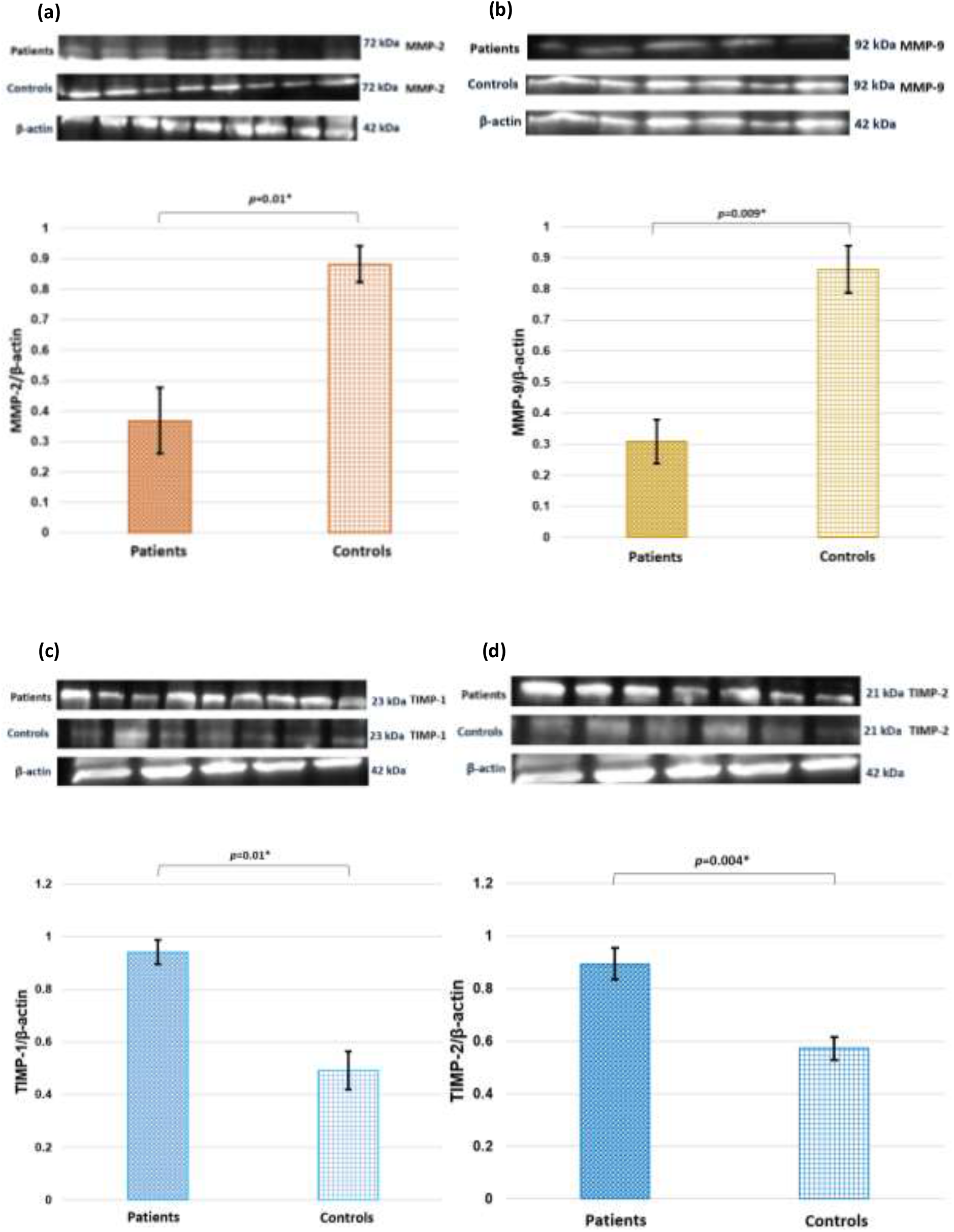
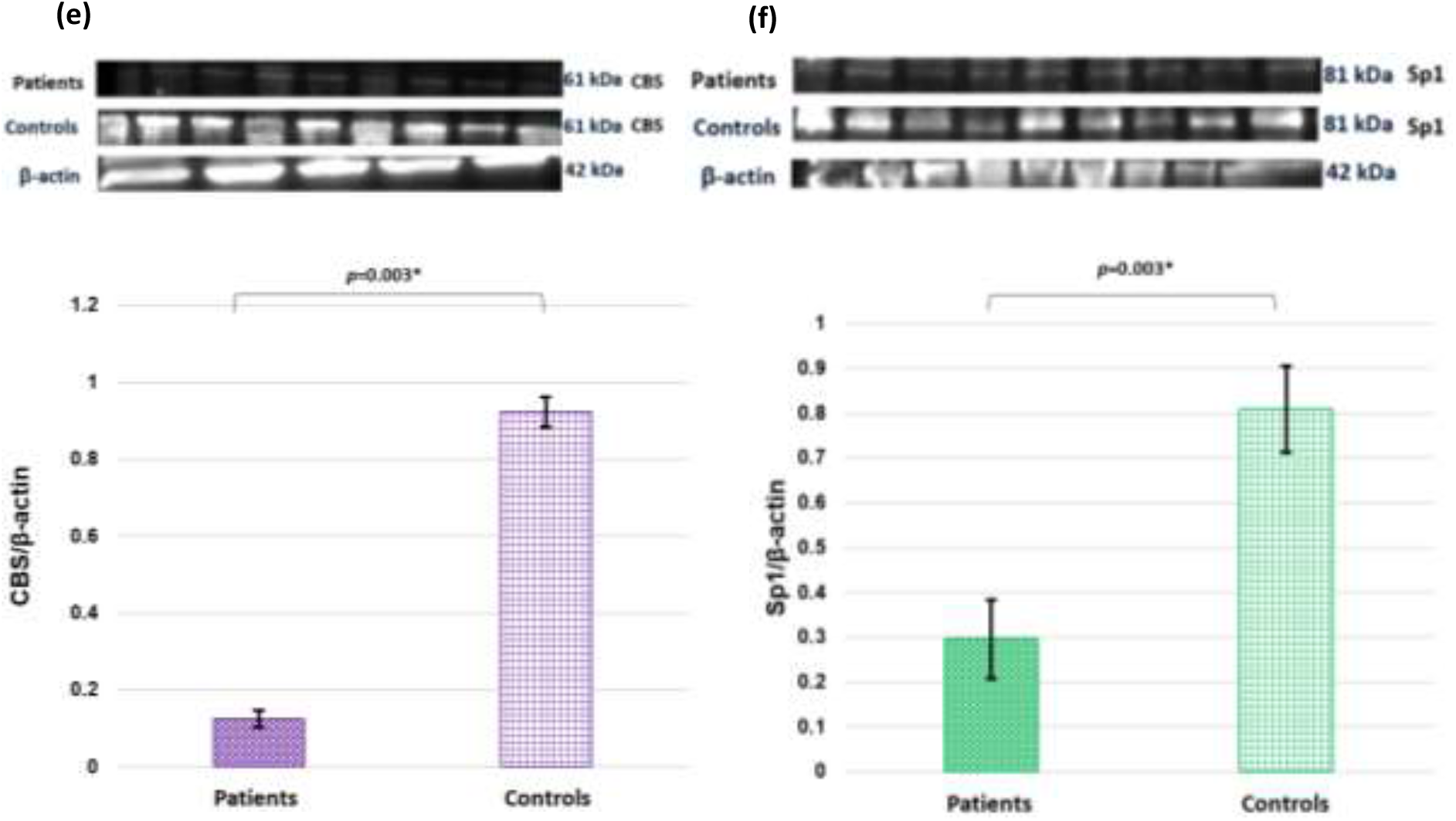
Representative images of immunoblot showing the protein expression of MMP 2 (a), MMP-9 (b), TIMP-1 (c), TIMP-2 (d), CBS (e) and Sp1 (f) in placental tissues of EOPE patients and normotensive, non-proteinuric controls. Bar diagrams represent the normalized values of MMPs 2, 9 (a,b), TIMPs 1, 2 (c,d), CBS (e) and Sp1 (f) with respect to β-actin (loading control). Data are presented as mean ± SEM. Statistical analysis was done using wilcoxon matched-pairs signed rank (MMPs 2, 9, TIMP-1, CBS, Sp1) and paired t (TIMP-2) tests, *p* values indicated on graphs.

**Figure 5:**
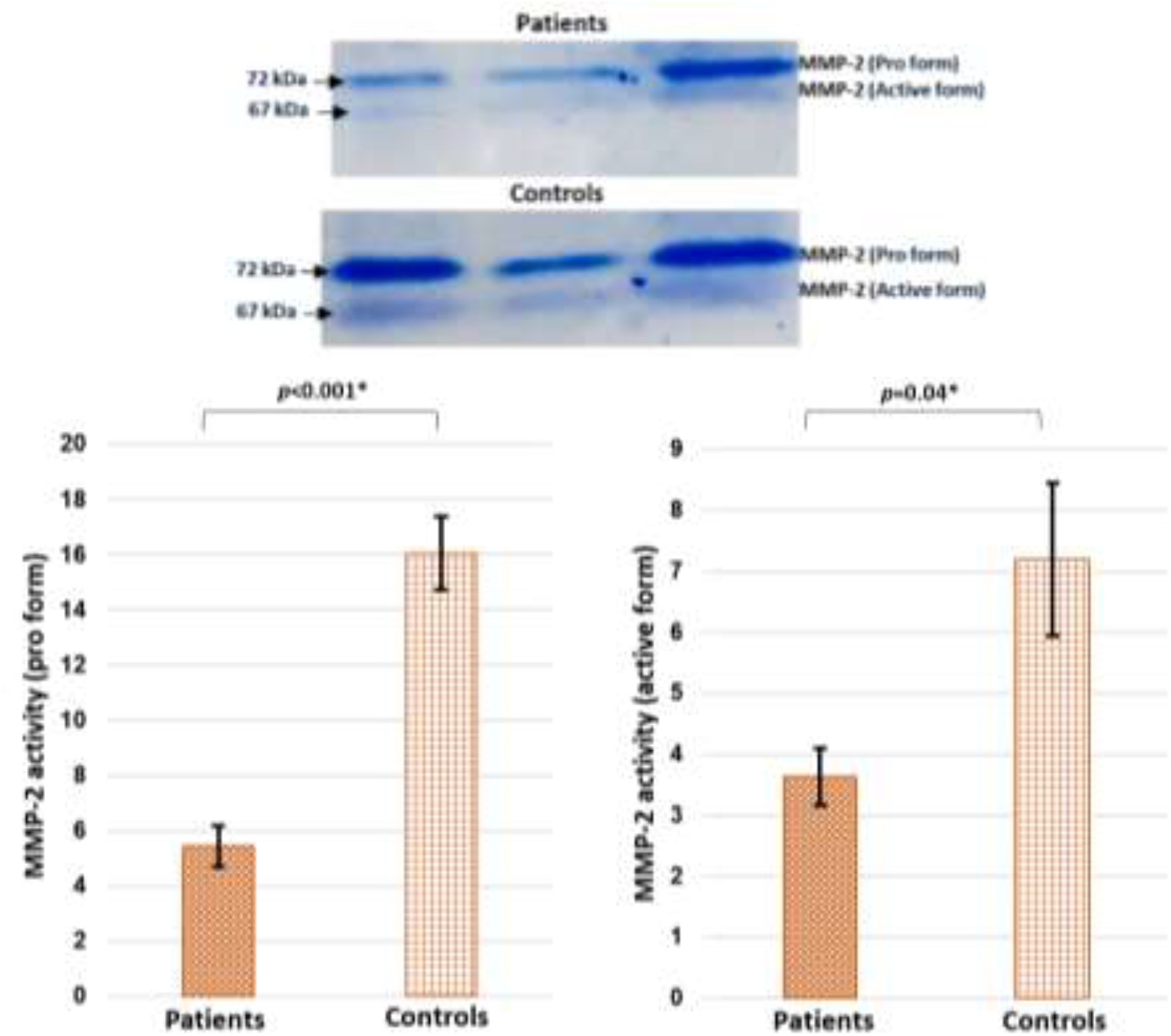
Representative zymograms depicting gelatinase activity on maternal side of placentae from early onset preeclamptic patients and normotensive, non-proteinuric controls. Comparison of Pro (72 KDa) and active (67 KDa) forms of MMP-2 between patients and controls was done by wilcoxon matched-pairs signed rank test. Data are expressed in the form of mean ± SEM, *p*<0.05 considered statistically significant.

**Figure 6:**
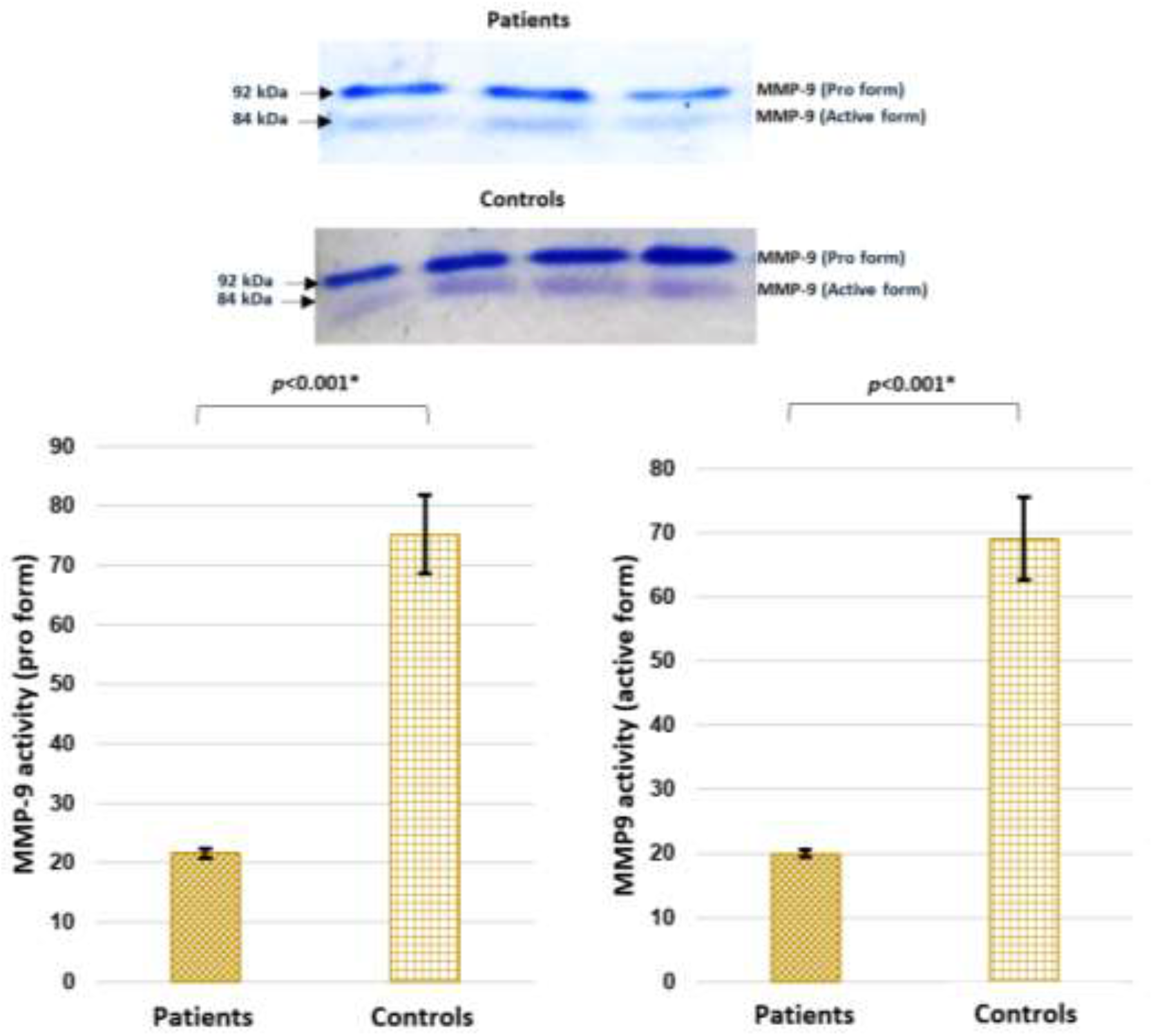
Representative zymograms depicting MMP-9 activity on maternal side of placentae from early onset preeclamptic patients and normotensive, non-proteinuric controls. Comparison of Pro (92 KDa) and active (84 KDa) forms of MMP-9 between preeclamptic patients and controls was done by wilcoxon matched-pairs signed rank test. Data are expressed in the form of mean ± SEM, *p*<0.05 was considered statistically significant.

### Significantly reduced expression of Cystathionine β-synthase (CBS) and Specificity protein 1 (Sp1) in the early onset preeclamptic patients

qRT-PCR data revealed significantly lower mRNA expression of CBS and Sp1 in placentae of early onset preeclamptic patients as compared to the normotensive, non-proteinuric controls [Figure 1]. IHC and IF staining demonstrated weaker expression of CBS and Sp1 in the syncytiotrophoblasts in the placentae from early onset PE patients as compared to control placentae [Figure 2,3]. Immunoblot data showed significantly lesser expression of CBS and Sp1 in the placentae from early onset PE patients as compared to normotensive, non-proteinuric control placentae [Figure 4].

### Significantly up regulated expression of precursor miR-22 and miR-22-3p in the early onset preeclamptic patients

qRT-PCR data revealed significantly up regulated miR-22 (precursor miR-22 and miR-22-3p) expression in the placentae from pregnancies complicated by early onset PE as compared to the normotensive, non-proteinuric controls [Figure 1].

## Discussion

Preeclampsia (PE), one of the most common pregnancy complications, is characterized by disturbed and inadequate remodelling of the maternal spiral arteries by invading trophoblast cells, thus reducing blood flow to the intervillous space^20^. Insufficient conversion of the spiral arteries into low-resistance, high-capacity vessels in early pregnancy leads to systemic hypertension and foetal hypoxia in later pregnancy as the foetus and placenta outgrow their blood supply, often observed in PE^21^. According to WHO, PE incidence is 7 times greater in developing countries compared to developed countries^22^. PE is classified into early (EOPE) and late onset and is widely accepted that these two entities have different aetiologies and should be regarded as different forms of disease^23^. EOPE is the most severe clinical variant of disease occurring in 5-20% of all cases of PE and is associated with impaired foetal growth, foetal pathology and uterine blood circulation, small size of the placenta, preterm delivery, neonatal morbidity and mortality^24^. LOPE is occurring in about 75-80% of all cases of preeclampsia, associated with maternal morbidity (metabolic syndrome, impaired glucose tolerance, obesity, dyslipidaemia, chronic hypertension) but normal birth weight and normal placental volume^24^. EOPE developments are associated with impaired trophoblast invasion, immune maladaptation and increased markers of endothelial dysfunction^24^.

To facilitate trophoblast cells invasion and adequate spiral artery remodelling, biochemical mediators such as matrix metalloproteinases (MMPs) are essential^25-27^. MMPs, also called matrixins, are a family of 17 zinc-dependent endopeptidases, which participate in many biological processes^28^. Specifically, MMP-2 and MMP-9 are involved in remodelling of placental and uterine arteries, and abnormal expression of these MMPs has been observed in hypertensive disorders of pregnancy^29,30^. In the present study, we have observed significantly down regulated gene and protein expression/levels of MMPs 2, 9 in the early onset preeclamptic patients as compared to normotensive, non-proteinuric controls by qRT-PCR, immunohistochemistry, immunofluorescence and western blot. Also, the gelatinolytic activity of MMP-2 and MMP-9 was found significantly lower in the early onset preeclamptic patients. Dang *et al*. in 2013 observed that MMP-2 and MMP-9 were abundantly expressed in tissues of normal pregnant rats by immunohistochemistry, western blots and gelatin gel zymography analysis^34^. Kolben *et al*. in 1996 and Shokry *et al*. in 2009 have observed reduced MMP-9 protein expression in preeclamptic placental tissues collected at delivery as compared to placentae from controls^35,36^. One previous study reported that preeclamptic umbilical cord tissues (artery and vein) had lower MMP-2 and MMP-9 levels than healthy tissues^37^. Gelatin gel zymography analysis revealed that zymogram wells loaded with preeclamptic amniotic fluid did not present any MMP-9 bands whereas the authors have determined pro-MMP-9 levels in normal amniotic fluid^38^.

MMP activity is regulated at the level of transcription, activation of latent forms, and inhibition by endogenous MMP inhibitors (tissue inhibitors of metalloproteinases; TIMPs)^31^. MMPs 2, 9 activity is regulated at different levels by interaction with TIMPs 2, 1^32, 33^. We, in the present study have observed significantly elevated mRNA and protein expression/levels of TIMPs 1, 2 in the placentae from early onset PE patients as compared to normotensive, non-proteinuric controls. TIMP-1 levels were found up regulated in preeclamptic patients as compared to normotensive pregnant women^39-41^. Western blot analysis revealed elevated TIMP-2 protein levels in the aminiotic fluid of women who subsequently develop preeclampsia^38^. Protein concentration of TIMP-2 was found raised in the preeclamptic patients as compared to healthy pregnant women^42^.

The molecular mechanism behind the aberrant expression of MMPs 2, 9 and their inhibitors (TIMPs 1, 2) in the pregnancies complicated by early onset preeclampsia is unexplored. Liu *et al*. in 2017 have studied the effects of exogenous hydrogen sulfide (NaHS) on the proliferation and invasion of human bladder cancer cells^43^. They have treated human bladder cancer EJ cells with different concentrations of exogenous NaHS (H_2_S donor) and they observed that exogenous NaHS could promote both the cell proliferation and invasion abilities^43^. They also found that exogenous NaHS treatment significantly increased the expression levels of MMP-2 and MMP-9 in EJ cells and speculated that exogenous H_2_S could promote cell proliferation and invasion by upregulating the expression of MMP-2 and MMP-9 in human bladder cancer EJ cells^43^. Therefore, we hypothesized that cystathionine β-synthase [hydrogen sulphide (H_2_S) producing enzyme] may influence the process of trophoblast invasion by affecting levels of MMPs 2, 9. We, in the present study have determined the expression of cystathionine β-synthase (CBS) at both transcription (qRT-PCR) and translation (IHC, IF, western blot) levels and we observed significantly up regulated mRNA and protein expression/levels of CBS likewise MMPs 2, 9 in the placental samples of normotensive, non-proteinuric controls; however, for early onset PE patients, CBS expression likewise MMPs 2, 9 was found to be significantly down regulated at both transcription and translation levels.f

To understand the regulation of cystathionine β-synthase gene, there is a previous study by Maclean *et al*. in which they proposed that the human CBS gene promoters -1a and -1b are expressed in a limited number of tissues and are co-ordinately regulated with proliferation through a redox-sensitive mechanism^17^. In their co-transfection studies in Drosophila SL2 cells, they indicated that both promoters (−1a and -1b) got transactivated by specificity protein 1 (Sp1) and specificity protein 3 (Sp3) but only the -1b promoter was subjected to a site-specific synergistic regulatory interaction between Sp1 and Sp3^17^. They proposed that Sp1 and Sp3 form a ternary complex with each other prior to binding the CBS -1b promoter region, as Sp1 binding has previously been shown to be capable of inducing conformational changes in DNA^44,45^, they speculated that one of the roles of Sp1 is to induce conformational changes in the CBS -1b promoter region that facilitate concomitant binding of Sp3 with a resultant synergistic increase in CBS promoter activity^17^. They also have observed that Sp1-deficient fibroblasts expressing both Sp3 and NF-Y were negative for CBS activity and transfection of these cells with a mammalian Sp1 expression construct induced high levels of CBS activity indicating that Sp1 has a critical and indispensable role in the regulation of CBS^17^. In the present study, we observed significantly up regulated mRNA and protein expression of Sp1 likewise CBS in the normotensive, non-proteinuric controls; however, for early onset PE patients, expression of Sp1 likewise CBS was found to be significantly down regulated at both transcription and translation levels.

Xia *et al*. in 2017 have identified Sp1, a novel, direct target of miR-22 using luciferase reporter assays^18^. They observed that miR-22 overexpression diminished but miR-22 knockdown increased Sp1 mRNA and protein expression in colorectal cancer cells^18^. Their observations provided strong evidences that miR-22 could partially inverse Sp1 induced proliferation, colony formation, migration and invasion in colorectal cancer cells and concluded that suppression of miR-22 by Sp1 activation is important for colorectal cancer cell growth and metastasis^18^. In the present study, we also have found inverse pattern of expression of Sp1 and miR-22 i.e., we have observed down regulated gene expression of Sp1 and up regulated for that of miR-22 (pre miR-22 and miR-22-3p) in the pregnancies complicated by early onset preeclampsia as compared to normotensive, non-proteinuric controls. Shao *et al*. in 2017 have reported enhanced miR-22 expression in early onset PE placentae as compared to their gestational age matched preterm labor controls^19^. They speculated that abnormally elevated levels of testosterone in the PE placentae may induce the expression of miR-22, which can directly inhibit estrogen receptor α levels and hamper E2/ERα signaling, leading to the downregulation in aromatase (CYP19A1) expression and subsequent inhibition of estradiol synthesis^19^. We, in our study have done qRT-PCR to determine the gene expression of pre miR-22 and miR-22-3p in the early onset PE patients and controls and observed significantly up regulated levels of miR-22 (pre miR-22 and miR-22-3p) in the early onset PE patients as compared to the normotensive, non-proteinuric controls.

## Summary

This is the very first study of its kind in which the mRNA and protein expression/levels of MMPs 2, 9, CBS and Sp1 were found to be significantly down regulated whereas up regulated for those of TIMPs 1, 2 in the early onset preeclamptic pregnancies as compared to their gestational and maternal age matched normotensive, non-proteinuric controls. We also have observed significant up regulation in the expression of miR-22 in the pregnancies complicated by EOPE as compared to controls. The aberrant functioning of these markers could possibly have contributed to the defective trophoblast invasion in the early onset preeclamptic patients in our study. The present study may be relevant for designing future screening programs in order to predict and to plan therapies for early onset preeclamptic patients.

## References

1. Portelli, M. and Baron, B. Clinical presentation of Preeclampsia and the diagnostic value of proteins and their methylation products as biomarkers in pregnant women with Preeclampsia and their newborns. Journal of Pregnancy., 2632637, pp. 23 (2018).

2. Sahay, A.S., Sundrani, D.P., Joshi, S.R. Regional changes of placental vascularization in preeclampsia: A review. IUBMB Life., 67, pp. 619–625 (2015).

3. Merchant, S.J. and Davidge, S.T. The role of matrix metalloproteinases in vascular function: Implications for normal pregnancy and pre-eclampsia. BJOG Int. J. Obst. Gynaecol., 111, pp. 931–939 (2004).

4. Myers, J.E., Merchant, S.J., Macleod, M., Mires, G.J., Baker, P.N., Davidge, S.T. MMP-2 levels are elevated in the plasma of women who subsequently develop preeclampsia. Hypertens. Pregnancy., 24, pp. 103–115 (2005).

5. American College of Obstetricians and Gynecologists and Task Force on Hypertension in Pregnancy, “Hypertension in pregnancy. Report of the American college of obstetricians and gynecologists’ task force on hypertension in pregnancy,” Obstetrics & Gynecology., 122 (5), pp. 1122–1131 (2013).

6. Tranquilli, A.L., Dekker, G., Magee, L. et al. The classification, diagnosis and management of the hypertensive disorders of pregnancy: a revised statement from the ISSHP. Pregnancy Hypertension: An International Journal of Women’s Cardiovascular Health., 4 (2), pp. 97–104 (2014).

7. Magee, L.A., Pels, A., Helewa, M., Rey, E. and von Dadelszen, P. Diagnosis, evaluation, and management of the hypertensive disorders of pregnancy. Pregnancy Hypertension: An International Journal of Women’s Cardiovascular Health., 4(2), pp. 105–145 (2014).

8. Tranquilli, A.L., Brown, M.A., Zeeman, G.G., Dekker, G. and Sibai, B.M. The definition of severe and early-onset preeclampsia. Statements from the international society for the study of hypertension in pregnancy (ISSHP). Pregnancy Hypertension: An International Journal of Women’s Cardiovascular Health., 3(1), pp. 44–47 (2013).

9. Poon, L.C. and Nicolaides, K.H. Early prediction of preeclampsia. Obstetrics and Gynecology International., 297397, pp. 11 (2014).

10. Paruk, F. and Moodley, J. Maternal and neonatal outcome in early- and late-onset pre-eclampsia. Seminars in Neonatology, 5(3), pp. 197–207 (2000).

11. Aksornphusitaphong, A. and Phupong, V. Risk factors of early and late onset pre-eclampsia. Journal of Obstetrics and Gynaecology Research., 39(3), pp. 627–631 (2013).

12. Stergiotou, I., Crispi, F., Valenzuela-Alcaraz, B., Bijnens, B. and Gratacos, E. Patterns of maternal vascular remodeling and responsiveness in early versus late onset preeclampsia. American Journal of Obstetrics and Gynecology., 209 (6), pp. 558.e1– 558.e14 (2013).

13. Sosa, S.E.Y., Flores-Pliego, A., Espejel-Nuñez, A., Medina-Bastidas, D., Vadillo-Ortega, F., Zaga-Clavellina, V. and Estrada-Gutierrez, G. New Insights into the role of matrix metalloproteinases in Preeclampsia. Int. J. Mol. Sci., 18, pp. 1448 (2017).

14. Zhen, Y. et al. Exogenous hydrogen sulfide exerts proliferation/anti-apoptosis/angiogenesis/migration effects via amplifying the activation of NF-kappaB pathway in PLC/PRF/5 hepatoma cells. Int J Oncol., 46, pp. 2194–2204 (2015).

15. Wu, D., Li, M., Tian, W., Wang, S., Cui, L., Li, H., Wang, H., Ji, A. and Li, Y. Hydrogen sulfide acts as a double-edged sword in human hepatocellular carcinoma cells through EGFR/ERK/MMP-2 and PTEN/AKT signaling pathways. Scientific Reports., 7(1), pp. 5134.

16. Sen, U., Sathnur, P.B., Kundu, S., Givvimani, S., Coley, D.M., Mishra, P.K., Qipshidze, N., Tyagi, N., Metreveli, N. & Tyagi, S.C. Increased endogenous H_2_S generation by CBS, CSE, and 3MST gene therapy improves ex vivo renovascular relaxation in hyperhomocysteinemia. Am J Physiol Cell Physiol., 303, pp. C41–C51 (2012).

17. Maclean, K.N., Kraus, E. & Kraus, J.P. The dominant role of Sp1 in regulating the cystathionine beta-synthase -1a and -1b promoters facilitates potential tissue-specific regulation by Kruppel-like Factors. The Journal of Biological Chemistry., 279(10), pp. 8558–8566 (2003).

18. Xia, S.S., Zhang, G.J., Liu, Z.L., Tian, H.P., He, Y., Meng, C.Y., Li, L.F., Wang, Z.W., Zhou, T. MicroRNA-22 suppresses the growth, migration and invasion of colorectal cancer cells through a Sp1 negative feedback loop. Oncotarget, 8 (22), pp: 36266–36278 (2017).

19. Shao, X., Liu, Y., Liu, M., Wang, Y., Yan, L., Wang, H., Ma, L., Li, Y.X., Zhao, Y., Wang, Y.L. Testosterone represses estrogen signaling by upregulating miR-22 A mechanism for imbalanced steroid hormone production in preeclampsia. Hypertension., 69, pp. 721–730 (2017).

20. Fisher, S.J. Why is placentation abnormal in preeclampsia? Am J Obstet Gynecol., 213(4): pp. S115–S122 (2015).

21. Ilekis, J.V., Tsilou, E., Fisher, S., Abrahams, V.M., Soares, M.J., Cross, J.C., Zamudio, S., Illsley, N.P., Myatt, L., Colvis, C., Costantine, M.M., Haas, D.M., Sadovsky, Y., Weiner, C., Rytting, E. and Bidwell, G. Placental origins of adverse pregnancy outcomes: potential molecular targets-an executive workshop summary of the Eunice Kennedy Shriver National Institute of Child Health and Human Development. Am J Obstet Gynecol., 215(1 Suppl), pp. S1–S46 (2016).

22. Kooffreh, M.E., Ekott, M., Ekpoudom, D.O. The prevalence of preeclampsia among pregnant women in the University of Calabar Teaching Hospital, Calabar. Saudi Journal for Health Sciences., 3(3), pp. 133–136 (2014).

23. Huppertz, B. Placental origins of preeclampsia: challenging the current hypothesis. Hypertension., 51(4), pp. 970–975 (2008).

24. Madazli, R., Yuksel, M.A., Imamoglu, M., Tuten, A., Oncul, M. et al. Comparison of clinical and perinatal outcomes in early and late onset preeclampsia. Arch Gynecol Obstet., 290(1), pp. 53–57 (2014).

25. Chen, J., Khalil, R.A. Matrix Metalloproteinases in normal pregnancy and preeclampsia. Prog Mol Biol Transl Sci., 148, pp. 87–165 (2017).

26. Zhu, J.Y., Pang, Z.J., Yu, Y.H. Regulation of trophoblast invasion: The role of Matrix Metalloproteinases. Rev Obstet Gynecol., 5(3-4), pp. e137–e143 (2012).

27. Burrows, T.D., King, A. and Lok, Y.W. Trophoblast migration during human placental implantation. Human Reproduction Update., 2(4), pp. 307–321 (1996).

28. Cui, N., Hu, M. and Khalil, R.A. Biochemical and Biological Attributes of Matrix Metalloproteinases. Prog Mol Biol Transl Sci., 147, pp. 1–73 (2017).

29. Isaka, K., Usuda, S., Ito, H., et al. Expression and activity of matrix metalloproteinase 2 and 9 in human trophoblasts. Placenta., 24, pp. 53–64 (2003).

30. Shimonovitz, S., Hurwitz, A., Dushnik, M., Anteby, E., Geva-Eldar, T., Yagel, S. Developmental regulation of the expression of 72 and 92 kd type IV collagenases in human trophoblasts: a possible mechanism for control of trophoblast invasion. Am J Obstet Gynecol., 171, pp. 832–8 (1994).

31. Lambert, E., Dasse, E., Haye, B., Petitfrere, E. TIMPs as multifacial proteins. Crit Rev Oncol Hematol., 49, pp. 187–98 (2004).

32. Isaka, K., Usuda, S., Ito, H., Sagawa, Y., Nakamura, H., Nishi, H., Suzuki, Y., Li, Y.F., Takayama, M. Expression and activity of matrix metalloproteinase 2 and 9 in human trophoblasts. Placenta., 24(1), pp. 53–64 (2003).

33. Van den Steen, P.E., Dubois, B., Nelissen, I., Rudd, P.M., Dwek, R.A., Opdenakker, G. Biochemistry and molecular biology of gelatinase B or matrix metalloproteinase-9 (MMP-9). Crit Rev Biochem Mol Biol., 37, pp. 375–536 (2002).

34. Dang, Y., Li, W., Tran, V., Khalil, R.A. EMMPRIN-mediated induction of uterine and vascular matrix metalloproteinases during pregnancy and in response to estrogen and progesterone. Biochemical pharmacology., 86(6), pp. 734–747 (2013).

35. Kolben, M., Lopens, A., Blaser, J. et al. Proteases and their inhibitors are indicative in gestational disease. Eur J Obstet Gynecol Reprod Biol., 68, pp. 59–65 (1996).

36. Shokry, M., Omran, O.M., Hassan, H.I., Elsedfy, G.O., Hussein, M.R. Expression of matrix metalloproteinases 2 and 9 in human trophoblasts of normal and preeclamptic placentas: preliminary findings. Exp Mol Pathol., 87, pp. 219–25 (2009).

37. Romanowicz, L., Galewska, Z. Extracellular Matrix Remodeling of the Umbilical Cord in Preeclampsia as a Risk Factor for Fetal Hypertension. J Pregnancy., 542695 (2011).

38. Lavee, M., Goldman, S., Daniel-Spiegel, E., Shalev, E. Matrix metalloproteinase-2 is elevated in midtrimester amniotic fluid prior to the development of preeclampsia. Reprod Biol Endocrinol., 7, pp. 85 (2009).

39. Palei, A.C., Sandrim, V.C., Cavalli, R.C., Tanus-Santos, J.E. Comparative assessment of matrix metalloproteinase (MMP)-2 and MMP-9, and their inhibitors, tissue inhibitors of metalloproteinase (TIMP)-1 and TIMP-2 in preeclampsia and gestational hypertension. Clin Biochem., 41, pp. 875–80 (2008).

40. Palei, A.C., Sandrim, V.C., Amaral, L.M. et al. Matrix metalloproteinase-9 polymorphisms affect plasma MMP-9 levels and antihypertensive therapy responsiveness in hypertensive disorders of pregnancy. Pharmacogenomics J., 12(6), pp. 489–98 (2011).

41. Montagnana, M., Lippi, G., Albiero, A., et al. Evaluation of metalloproteinases 2 and 9 and their inhibitors in physiologic and pre-eclamptic pregnancy. J Clin Lab Anal., 23, pp. 88–92 (2009).

42. Palei, A.C., Sandrim, V.C., Amaral, L.M. et al. Association between matrix metalloproteinase (MMP)-2 polymorphisms and MMP-2 levels in hypertensive disorders of pregnancy. Exp Mol Pathol., 92, pp. 217–21 (2012).

43. Liu, H., Chang, J., Zhao, Z., Li, Y., Hou, J. Effects of exogenous hydrogen sulfide on the proliferation and invasion of human Bladder cancer cells. J Can Res Ther., 13 (5), pp. 829–832 (2017).

44. Su, W., Jackson, S., Tjian, R. and Echols, H. DNA looping between sites for transcriptional activation: self-association of DNA-bound Sp1. Genes Dev., 5, pp. 820– 826 (1991).

45. Ikeda, K., Nagano, K. and Kawakami, K. Anomalous interaction of Sp1 and specific binding of an E-box-binding protein with the regulatory elements of the Na,K-ATPase α2 subunit gene promoter. Eur. J. Biochem., 218, pp. 195–204 (1993).

